# Lack of ADAP1/Centaurin-α1 Ameliorates Cognitive Impairment and Neuropathological Hallmarks in Mouse Alzheimer Model

**DOI:** 10.1101/2025.01.05.630617

**Authors:** Erzsebet M. Szatmari, Corey Moran, Sarah J. Cohen, Denys Bashtovyy, Amanda Jacob, Wyatt Bunner, Mary Phipps, Joan Cristino Lora, Robert W. Stackman, Ryohei Yasuda

## Abstract

ArfGAP, with dual PH domain-containing protein 1/Centaurin-α1 (ADAP1CentA1), is a brain-enriched and highly conserved Arf6 GTPase-activating and Ras-anchoring protein. ADAP1 is involved in dendritic outgrowth and arborization, synaptogenesis, and axonal polarization by regulating the dynamics of the actin cytoskeleton. An increased level of ADAP1 and its association with amyloid plaques in the human Alzheimer’s disease (AD) brain suggest a role for this protein in AD progression. To understand the role of ADAP1/CentA1 in neurodegeneration, we crossbred CentA1 KO mice with the hAPP-J20 mouse model of AD (J20 x CentA1 KO). We then evaluated the gene expression profile and the behavioral and neuropathological hallmarks of AD to determine the impact of eliminating ADAP1/CentA1 expression on AD-related phenotypes. Spatial memory assessed by the Morris Water Maze test showed significant impairment in J20 mice, which was rescued by the deletion of CentA1. Neuropathological hallmarks of AD, such as deposits of amyloid plaques and neuroinflammation, were significantly reduced in the AD model mice with CentA1 KO background. To identify potential mediators of AD phenotype rescue, we analyzed differentially expressed genes (DEGs) between genotypes, by employing transcriptome profiling with Nanostring nCounter Neuropathology and Neuroinflammation panels. We found significant upregulation of genes associated with apoptosis and gliosis in the brain of J20 mice. However, many of these genes, including the pro-apoptotic gene, *Bid*, were restored in the brains of J20 x CentA1 KO mice compared to J20 mice. In summary, our data indicate that CentA1 is required for the progression of AD phenotypes and that targeting CentA1 signaling at mitochondria might have therapeutic potential for AD prevention or treatment.

**SIGNIFICANCE STATEMENT:** ADAP1/Centaurin-α1 (CentA1) is highly enriched in the brain and increased CentA1 level has been linked to Alzheimer’s disease (AD). However, the precise role of ADAP1 in the pathogenesis of AD is poorly understood. We found that genetic deletion of CentA1 in the AD model mice rescues the pathological hallmarks of AD, including loss of dendritic spines in the hippocampus, amyloid plaque deposition, neuroinflammation and spatial memory deficits. Transcriptome analysis of forebrain samples using NanoString nCounter panels for 880 genes, identified the pro-apoptotic protein, Bid, with significantly reduced expression in the J20 mice on ADAP1 KO background. These findings point towards a role of ADAP1 in the activation of the mitochondrial pathways of death associated with the progression of neurodegeneration.

## INTRODUCTION

Alzheimer’s disease (AD) is a pervasive neurodegenerative condition characterized by progressive decline of memory and cognitive function^1–3^. Extracellular amyloid plaque depositions, formation of intracellular neurofibrillary tangles, widespread neuroinflammation, neuronal cell death, and alterations in brain morphology are the major pathologies of AD^4–6^. However, the molecular complexity underlying these pathological transformations is poorly understood. Early in AD, soluble Aβ oligomer-induced loss of dendritic spines, aberrant synaptic remodeling, and abnormal synaptic transmission cause deficits in the activity of the networks mediating cognitive functions and learning and memory^7–10^. Several molecular signaling mechanisms have been implicated in the Aβ-triggered synaptic dysfunction, including eIK2α kinases, NMDA receptor signaling, the Ras-ERK pathway, CAMKK2-AMPK kinase pathway, GSK3β and the ADAP1/Centaurin-α1-Ras-Elk1 signaling at mitochondria^11–15^. ADAP1/CentA1 (ArfGAP with dual PH domain-containing protein 1/Centaurin-α1) is highly expressed in brain areas involved in AD, including the hippocampus^16–18^. In neurons, ADAP1 is expressed in the axon^19,20^, the dendrite, dendritic spines, and the nucleus ^21,22^, and localized to the plasma membrane and the mitochondria ^23^.

Structurally, ADAP1 is a multi-domain protein with an ArfGAP domain, which targets Arf6 in vivo^24,25^, and two PH domains. ADAP1 interacts with several signaling molecules, including PI3K, cytoskeletal, nuclear and mitochondrial proteins and cytosolic kinases, providing a scaffolding platform for these proteins^23^. Additionally, ADAP1 interacts with Ras and facilitates the activation of the Ras-ERK1/2 pathway^11,26^.

Several studies, including ours, suggested that ADAP1/CentA1 is involved in AD. In the postmortem human AD brain, the level of CentA1 increases, particularly around neuritic plaques^23,27,28^. Using cellular models of AD, we previously reported that in cultured neurons and hippocampal slices, Aβ-induced dendritic spine loss and aberrant spine structural plasticity are significantly suppressed if CentA1 is downregulated via shRNA^11^. In addition, to understand the role of ADAP1/CentA1 in health and disease, we created ADAP1/CentA1 global knockout (KO) mice. We found that deletion of ADAP1/CentA1 leads to enhanced dendritic spine density and structural LTP in the hippocampus and improved performance in a hippocampal-dependent spatial memory task^29^.

In this study, to verify the involvement of ADAP1/CentA1 in the molecular pathogenesis of AD, we evaluated AD-related phenotypes in the hAPP-J20 mice on an ADAP1/CentA1 KO background. The hAPP-J20 mouse model overexpresses human APP with the Swedish and Indiana mutations. These mice simulate several hallmarks of sporadic and autosomal dominant AD, including neuroinflammation, cerebral Aβ plaque burden, synaptic dysfunction, spontaneous epileptic activity, and deficits in spatial memory and learning^30–33^. By using conventional histological analysis, biochemical studies, gene expression profiling and behavioral assay, we found that the removal of ADAP1/CentA1 rescues spatial memory deficits in AD mice, while ameliorating the histopathological hallmarks of neurodegeneration, including dendritic spine elimination, amyloid plaque deposition and inflammatory elevation. Using gene expression profiling, we explored the translational potential of ADAP1-dependent rescue of AD-phenotypes. Analysis of differentially expressed genes (DEG) and pathways revealed that the fundamental themes of neurodegeneration most affected by lack of ADAP1/CentA1 are those associated with aging, neuroplasticity, and development; neurotransmission; structural integrity and compartmentalization and metabolism. We conclude that ADAP1/CentA1 signaling represents a target for therapeutic approaches to treat AD.

## MATERIALS AND METHODS

### Animals

All mice were housed in the Animal Resource Facility of Max Planck Florida Institute for Neuroscience, compliant with the US National Institutes of Health Guide for Care and Use of Laboratory Animals.

ADAP1/Centaurin-α1 KO mice were generated as previously described ^29^ and then crossbred with the J20 mouse model of AD. J20 mice ^30^ were in-house breed by crossing heterozygous and WT (C57BL/6) mice. J20 mice were heterozygous for transgene. Genotyping was performed before experiments by PCR of genomic DNA extracted from ear snip material (Transnetyx). At the end of the experiment, genotype was reconfirmed by western blotting of brain samples. Mice were group-housed (2-4 mice/cage) and littermates were randomly distributed over different cages. Non-transgenic littermates referred to as wildtype (WT) mice were used as controls. Only male mice were used for this study. The experimental groups were J20, J20 crossed with CentA1/ADAP1 KO (J20 x KO), and controls were wild type littermates (WT), ensuring that the mice were from the same parents, living in the same cage and uniformly affected by environmental factors.

### Immunofluorescence staining

Adult male mice (4-7 months old) were deeply anesthetized with Ketamine/Xylazine cocktail until lack of response to toe pinch was recorded, then perfused transcardially with saline followed by perfusion with 4% paraformaldehyde in 0.1M phosphate buffer (PB). Brains were removed and post fixed overnight in the same fixative at 4°C. Coronal sections were cut at 50 µm on a Leica vibratome and collected in ice cold 0.1M PB. After a brief rinse with 0.1M PB, the free-floating sections were rinsed in 0.1M PB then incubated for 30 minute in blocking buffer (0.3% Triton and 0.5% normal goat serum in 0.1M PB). Sections were reacted overnight with anti-NeuN antibody (ABN78, rabbit polyclonal, Millipore, Temecula, CA, USA) diluted 1:1000 in blocking buffer. Sections were washed in 0.1M PB and then incubated for 2 hours with Alexa 488-conjugated secondary antibodies (A-11008, Life Technologies) diluted 1:500 in blocking buffer. Sections were rinsed in 0.1M PB and the nuclei were stained with Hoechst (1:10000, H3570, Life Technologies) for 10 minutes. Sections were rinsed again in 0.1M PB and mounted on Superfrost plus slides (Thermo Fisher) using Fluoromount-G. We imaged four sections from four mice for both genotypes using a Zeiss LSM 710 confocal microscope.

### SDS-PAGE and immunoblotting

Hippocampi were extracted with T-PER protein extraction buffer (Pierce) supplemented protease and phosphatase inhibitors (Roche). The lysates were centrifuged at 15000 g for 15 minutes at 4°C and the supernatants were used for further analysis. Samples were prepared for standard SDS-PAGE and separated on 4-20% gradient acrylamide gel (Mini-PROTEAN TGX precast gels, Bio-Rad), then transferred onto 0.45 µm pore size PVDF membranes (Millipore) using semi-dry immunoblotting.

Membranes were blocked with 5% nonfat milk in TBS-T (Tris Buffered Saline with 0.2% Tween-20) for 1 hour at room temperature, then incubated overnight at 4°C with primary antibodies diluted in 5% BSA in TBS-T. The following commercially available antibodies were used: mouse anti-β -amyloid (clone 6E10, Covance/Biolegend; 1:1000); goat anti-Centaurin-1 (Abcam; 1:500); rat anti-BID (R&D Systems; 1:500) and mouse anti-β-actin (Sigma, 1:1000). Membranes were washed 3 times for 15 minutes in TBS-T, followed by incubation for 2 hours at room temperature with HRP-conjugated donkey anti-goat; goat anti-rat; or rabbit anti-mouse secondary antibodies (Bio-Rad), diluted 1:2000 in 5% nonfat milk in TBS-T. Membranes were washed 3 times for 15 minutes in TBS-T, then incubated with Pierce ECL Plus western blotting substrate to detect western blotted proteins. We used the Bio-Rad Chemidoc imaging system to visualize protein bands. ImageJ software was used for western blot quantification.

### Morris Water Maze test

Spatial learning and memory were evaluated by the Morris Water Maze (MWM) test. Briefly, for 3 days before testing, mice were handled and habituated to the testing facility. One hour before testing, the mice were moved to a waiting room with light level set to that of the testing apparatus to ensure adjusting. The water maze consisted of a 1.4 m diameter white tank with a 10 cm diameter platform submerged approximately 1 cm below the surface of the water. The temperature of the water was kept at 22-24°C. The water was made opaque using non-toxic white washable paint to make the platform invisible during trials. Visual cues were placed in the testing room around the tank for spatial reference. Prior to water maze training, mice received a visual platform test where the spatial cues were removed, and the platform was elevated above the surface of the water and marked with a cue, so it could clearly be discerned. Mice were given four trials, and the platform location was varied over trials. This served to verify the visual ability of the mice and to ensure that the mice had no motor deficits that would affect their ability to swim to the platform. For the hidden platform test, mice were given four acquisition trials per day for eight consecutive days. The start location was varied for each trial, and the mice were allotted 60 seconds to find the platform. Mice were left on the platform for 15 seconds before removing them from the water maze. If a mouse did not find the platform within 60 seconds, it was placed on or guided to the platform and kept there for 15 consecutive seconds. Mice were dried after each trial and placed into cages located atop heating pads to prevent hypothermia. Daily acquisition trials were averaged for analysis. On day 9 mice were given a probe test during which the platform was not present. Activity and performance were tracked using EthoVision XT (Noldus Information Technology, Leesburg, VA). Total time spent in each quadrant, total number of entries into the target quadrant, total number of platform crossings, latency to first platform crossing, number of trials to criterion for reaching platform center (20 sec) and average distance to the platform center were recorded. All experiments were performed in a double-blinded manner.

### Golgi-Cox staining and analysis of dendritic spine density

We used a commercially available Golgi-Cox staining system, the FD Rapid Golgi Staining Kit. Briefly, mice were deeply anesthetized with Ketamine/Xylazine cocktail until lack of response to toe pinch was recorded. Brain samples were collected and rinsed with Milli-Q water. Impregnation was performed following the manufacturer’s instructions. Further processing of samples was done at FD Neurotechnologies. Images of pyramidal neurons from the CA1 area of the hippocampus were collected on 100 µm thick slices using a Zeiss 780 confocal laser-scanning microscope. To detect Golgi-Cox staining the microscope was set up in transmission mode and 488 nm wavelength laser was used. Confocal images were obtained using a Plan-Neofluar 63x water (1.3 numerical aperture) objective. Each frame was acquired eight times and then averaged to obtain noise-free images. Spine density was determined on secondary apical dendrites of CA1 pyramidal neurons in SLM and SR regions of the hippocampus. The number of spines/100 µm was calculated using ImageJ software.

### Immunohistochemistry staining

For IHC studies, mice were deeply anesthetized with Ketamine/Xylazine cocktail until lack of response to toe pinch was recorded. Mice were transcardially perfused with 50 mL PBS and 50 mL 4% paraformaldehyde (PFA) and the brain was harvested and postfixed in 4% PFA for at least 24 hours. Paraffin embedded sagittal brain sections (5µm thickness) were mounted on Superfrost Plus slides (Fisherbrand). Sections were deparaffinized, then either pretreated with 70% Formic Acid (6E10, Covance/Biolegend) or heat retrieved in a BioCare Decloaker (anti-GFAP; Abcam, anti-Iba1, Abcam). For blocking we used Rodent Block M (BioCare Medical, Concord, CA) for 30 minutes at room temperature. Samples were incubated overnight at 4°C with primary antibody diluted in Van Gogh Diluent (BioCare Medical, Concord, CA): 1:4000 (6E10, Covance), 1:200 (Iba1, ab12267, Abcam) and 1:1000 (anti-GFAP, Abcam). Sections were rinsed with TBS-T and incubated with secondary HRP-Polymers (Mouse-on-Mouse, Rabbit-on-Rodent or Goat-on-Rodent (BioCare Medical, Concord, CA) for 30 minutes at room temperature. After several TBS-T rinses the samples were reacted with Betazoid DAB (BioCare Medical, Concord, CA), followed by TBS washes, then stained with Hematoxylin, dehydrated and coverslipped with Leica Micromount mounting medium.

### Immunohistochemical imaging and image processing

Immunohistochemically stained sections were captured using a BZ-X800 digital microscope (Keyence; Chicago, IL, USA) and analyzed using the Aperio ImageScope program as previously described ^34^. Amyloid plaque burden and neuroinflammation in the forebrain (cortex and hippocampus) were calculated using the Positive Pixel Count program within the Aperio ImageScope software. Four sections per brain (n = 3-4 mice/group) that were cut at 30 mm apart, were analyzed and averaged for each mouse by an investigator blinded to genotype. The final images and layouts were created using Creative Cloud Photoshop and Illustrator (Adobe, San Jose, CA).

### RNA extraction and analysis

RNA was isolated from frozen mouse cortical tissue. Isolated RNA was purified using the Qiagen RNeasy kit following the manufacturer’s instructions. The concentration of RNA was determined using Nanodrop (Thermo Fisher) and the quality of RNA was confirmed using the ECU Genomic Core’s Bioanalyzer system with RNA pico chips.

### Gene expression analysis

Gene expression analysis was performed using NanoString Mouse Neuropathology and Neuroinflammation gene expression panel (NanoString Technologies Inc., Seattle, WA, USA). Briefly, RNA isolated from fresh-frozen cortical tissue was used for experiments. 100 ng of total RNA was hybridized with reporter and capture probes for nCounter Gene Expression code sets (Neuropathology and Neuroinflammation) according to the instructions from manufacturer (NanoString Technologies Inc., Seattle, WA, USA). Using the NanoString nSolver Analysis system, data were normalized to spiked positive controls and housekeeping genes. Transcript counts less than the mean of the negative control transcripts plus 2STDEV for each sample were considered background. RNA isolated from 3-4 mice/genotype was used for Nanostring mRNA analysis.

### Quantitative RT-PCR (qRT-PCR) analysis

Complementary DNA (cDNA) was synthesized with the High-Capacity cDNA Reverse Transcription Kit (Applied Biosystems) following manufacturer’s instructions. Quantitative RT-PCR was performed (POWER SYBR GREEN PCR Master Mix; Applied Biosystems) in triplicate using the Applied Biosystems ViiA 7 system. mRNA gene expression was normalized to mouse 18s ribosomal RNA, with expression fold change calculated using the 2^−^ΔΔCt method. Commercially available Bid Mouse qPCR Primer Pair (NM_007544) was used for qPCR (Origene; catalog number MP201328):

### Experimental Design and Statistical Analysis

For all experiments all genotypes were processed in parallel. Specific sexes (males only) were used for behavioral studies, immunofluorescence and immunohistology studies. GraphPad Prism (version 10 for Windows, GraphPad Software, San Diego, California USA, www.graphpad.com) was used for majority of statistical analysis. Student’s t test was used to compare two independent datasets. For multiple comparisons, we used one-way ANOVA followed by Tukey’s multiple-comparisons test. Differences between genotypes or samples were considered significant at p<0.05. Data are reported as mean ± S.E.M, unless otherwise stated.

## RESULTS

### Normal gross brain morphology of J20 and J20 x CentA1 KO mice

To assess whether ADAP1/CentA1 deletion affects gross brain structure in AD model mice, we compared the overall brain morphology between J20, J20 x KO and their non-transgenic (WT) littermates. NeuN immunostaining of coronal sections was indistinguishable between genotypes (**FIG. 1A**). Lack of CentA1 protein and expression of the hAPP transgene were verified by immunoblotting in the hippocampus of 6- (**FIG. 1B**) and 12-14-months old mice (**FIG. 1C**).

**FIG. 1:**
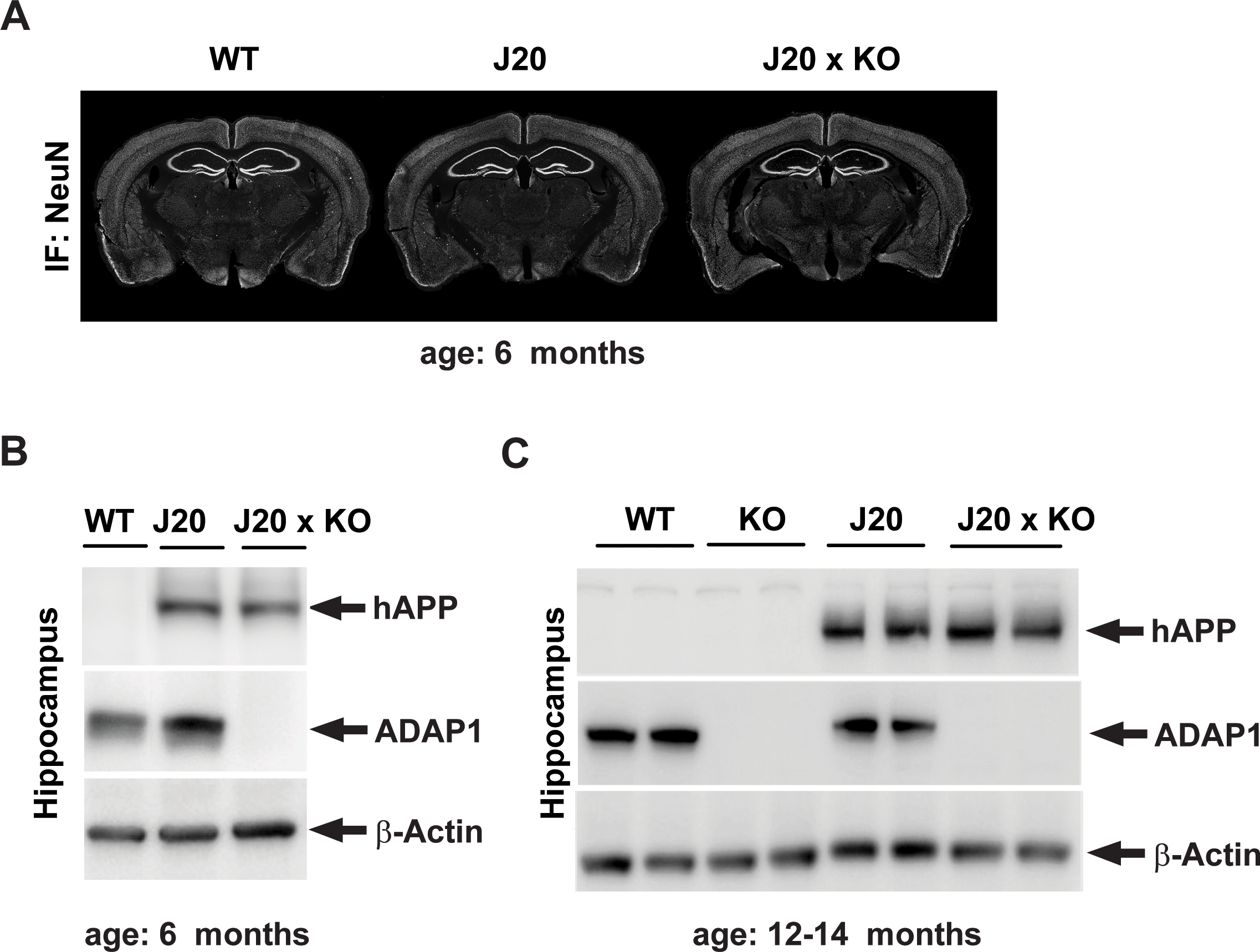
Generation and validation of hAPP-J20 x CentA1 KO mice. ***A***, NeuN immunohistochemistry of coronal sections from 6 months old hAPP-J20 (J20); hAPP-J20 x CentA1 KO (J20 x KO) and non-transgenic littermate (WT) mice show normal brain morphology in the J20 x KO mice. ***B***, Immunoblots show validation of J20 x KO mice: the presence of hAPP and the complete lack of CentA1 protein in the hippocampus of 6 months old J20 x KO mice. β-Actin was used as loading control. ***C***, Immunoblots show validation of J20 x KO mice: the presence of hAPP and the complete lack of CentA1 protein in the hippocampus of 12 months old J20 x KO mice. β-Actin was used as loading control. The number of mice was 2-3/genotype.

### Lack of ADAP1/CentA1 rescues spatial learning and memory deficit in the J20 AD model mice

We next assessed whether the deletion of CenA1 influences the behavioral phenotypes of the J20 mice through the Morris Water Maze (MWM) task to evaluate hippocampus-dependent spatial learning and memory. We trained the mice using four trials/day for 8 days, with an inter-trial interval of 20-30 minutes. As expected, a two-way repeated measures (genotype, trial block) ANOVA revealed the distance traveled to the target location decreased with training for all genotypes (F_(3,637)_ = 12.27, *p* = 0.001). Holm-Sidak post hoc analyses indicated that the J20 animals had a deficit in learning acquisition, that reached significance on day 4 of training (compared to WT: *t*_(44)_ = 3.16, *p* = 0.001; Average distance traveled for day 4 training, mean ± S.E.M; WT: 138.37 ± 5.95 cm, n = 26; KO: 138.07 ± 6.19 cm, n = 24; J20: 177.08 ± 6.07 cm, n = 25; J20 x KO: 148.31 ± 6.78 cm, n = 20) (**FIG. 2A**). Another parameter to analyze learning acquisition is the number of trials to reach a predefined group criterion of performance; which was set to 20 seconds of escape latency ^35^. A one-way (genotype) ANOVA on training trials needed to reach criterion revealed a significant genotype effect (F_(3, 91)_ = 6.37, *p* = 0.001). Holm-Sidak post hoc analyses indicated that it took two days longer for the J20 mice to reach this predefined criterion (compared to WT: *t*_(49_) = 3.18, *p* = 0.01), while lack of ADAP1/CentA1 rescued this deficit (compared to J20: *t*_(20)_ = 3.30, *p* = 0.001; average number of trials to reach criterion, mean ± S.E.M; WT: 11.42 ± 0.83 trials, n = 26; KO: 10.83 ± 1.21 trials, n = 24; J20: 17.80 ± 1.58 trials, n = 25; J20 x KO: 11.50 ± 1.40 trials, n = 20) (**FIG. 2B)**. Once learning criteria were met, however, all genotypes showed similar performance on the memory probe test administered on day 9 (one-way (genotype) ANOVA on the percent of crossings into the target search zone: F*_(3, 91)_* = 0.66, *p* = *n.s.*) (**FIG. 2C)**. These results suggest that J20 animals have a deficit in spatial learning, that is partially rescued by the removal of ADAP1/CentA1.

**FIG. 2:**
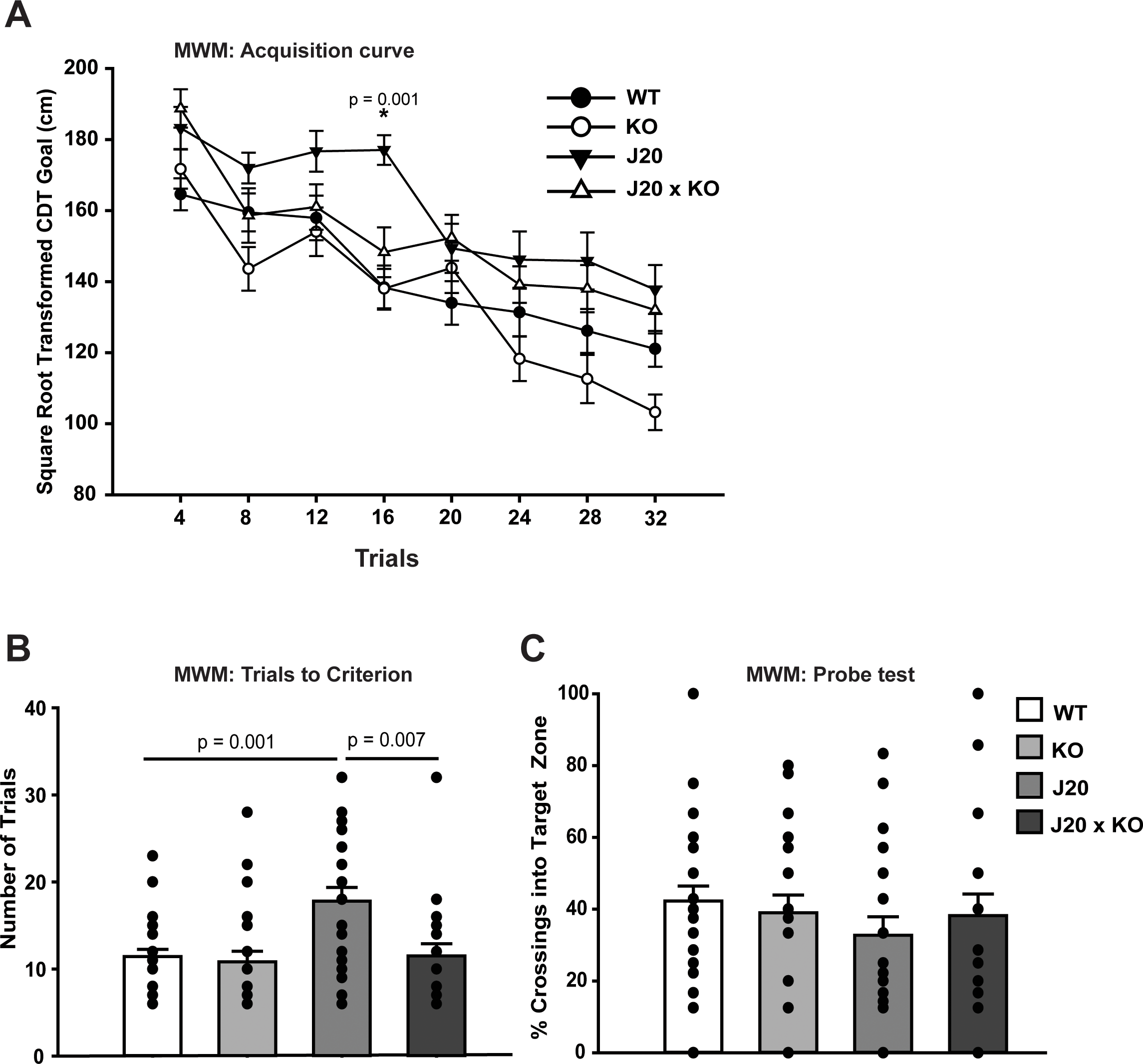
Lack of CentA1 protein rescues spatial memory deficit in the hAPP-J20 mice. ***A***, Acquisition curve of hippocampal-dependent spatial learning and memory evaluated by the MWM test. The distance traveled to reach the target escape platform decreased across training days for all genotypes. However, the hAPP-J20 mice demonstrated a significant deficit in learning acquisition by day 4 of training compared to J20 x KO mice (p = 0.001). ***B***, The number of trials needed to reach the predefined criterion of performance (escape latency maximum of 20 sec) was significantly higher for the J20 mice compared to WT (p = 0.001) and J20 x KO mice (p = 0.007), indicating rescue of learning deficit by the lack of ADAP1/CentA1. ***C***, All genotypes performed similarly on the memory probe test administered on day 9. The number of mice was WT: n = 26; KO: n = 24; J20: n = 25; J20 x KO: n = 20) XX-YY/genotype. **SARAH, CAN YOU HELP HERE?).** All behavior data are presented as mean ± SEM.

### Deletion of ADAP1/CentA1 rescues dendritic spines in the hippocampus of hAPP-J20 mice

Loss of dendritic spines and synapses are early pathological hallmarks of AD, that precedes plaque deposits in the AD brain ^36,37^.Therefore we analyzed the effect of the deletion of ADAP1/CentA1 on the loss of dendritic spines in the hippocampus CA1 neurons of J20 mice. We counted the number of dendritic spines in the stratum lacunosum moleculare (SLM) and the stratum radiatum (SR) of the hippocampus of J20 mice and their J20 x KO littermates (**FIG. 3)**. Five neurons/animal were included in analysis (n=4-6 mice/genotype).We found that spine density was significantly reduced in the SLM of J20 mice compared to WT, while ADAP1/CentA1 KO rescued this effect (**FIG. 3A;** number of spines/100 µm; WT= 107.4 ±1.28; J20 = 84.05±1.88; J20 x KO= 93.6 ±2.23; WT vs. APP: p=0.0001; WT vs. APP x KO: p= 0.0003; APP vs. APP x KO: p=0.01; S.E.M; one-way ANOVA followed by Tukey’s multiple comparisons test). Interestingly, there is a small but statistically significant increase in spine number in the SR area in J20 mice compared to WT, but J20 x KO showed normal spine number (**FIG. 3B;** number of spines/100 µm; WT= 93.25 ±3.83; J20 = 109.5±4.52; J20 x KO= 105.3 ±2.6; WT vs. J20: p=0.03; WT vs. J20 x KO: p= 0.1; J20 vs. J20 x KO: p=0.7; S.E.M; one-way ANOVA followed by Tukey’s multiple comparisons test).

**FIG. 3:**
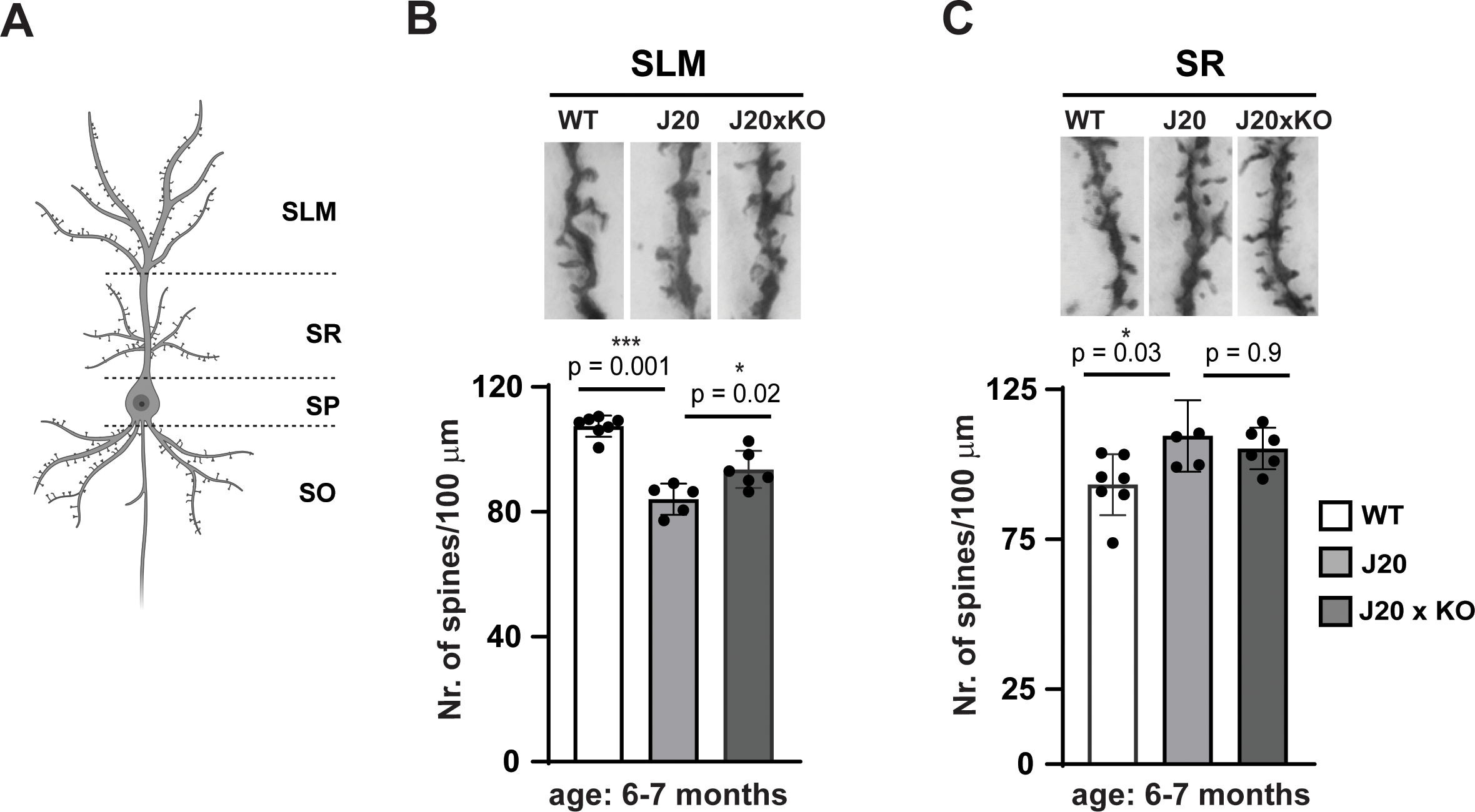
Lack of ADAP1/CentA1 rescues dendritic spine density in the hippocampus of J20 mice. ***A***, Graph shows significantly reduced dendritic spine density in the hippocampus lacunosum moleculare (SLM) neurons of J20 mice compared to WT litter mates (p=0.001). Lack of ADAP1/CentA1 prevented the loss of dendritic spines in the SLM of J20 mice on ADAP1/CentA1 KO background (p=0.02). ***B***, Dendritic spine density in the stratum radiatum (SR) of the hippocampus in J20 mice is significantly higher compared to WT littermates (p=0.03), while ADAP1/CentA1 KO does not rescue this effect (p=0.9). The number of mice was 4-5/genotype and 5 neurons/animal were analyzed. All data are presented as mean ± SEM.

### Deletion of ADAP1/CentA1 reduces amyloid plaque deposition in the hippocampus of hAPP-J20 mice

We analyzed the effect of ADAP1/CentA1 KO on the amyloid deposition in the hippocampus and the cortex of J20 mice and their J20 x KO littermates (**FIG. 4)**. Quantification of Aβ plaque immunoreactivity in the hippocampus (**FIG. 4A and B)** showed a significant reduction in plaque burden in the J20 mice lacking ADAP1/CentA1 (J20: 0.69±0.34%; n=4; J20 x KO: 0.43±0.21%: n=4: p=0.016; S.E.M; one-way ANOVA followed by Tukey’s multiple comparisons test). Amyloid deposition in the cortex showed no difference between J20 and J20 x KO mice (**FIG4. C)** (J20: 0.69±0.34%; n=4; J20 x KO: 0.49±0.24%: n=4: p=0.41; S.E.M; unpaired t test with Welch’s correction).

**FIG. 4:**
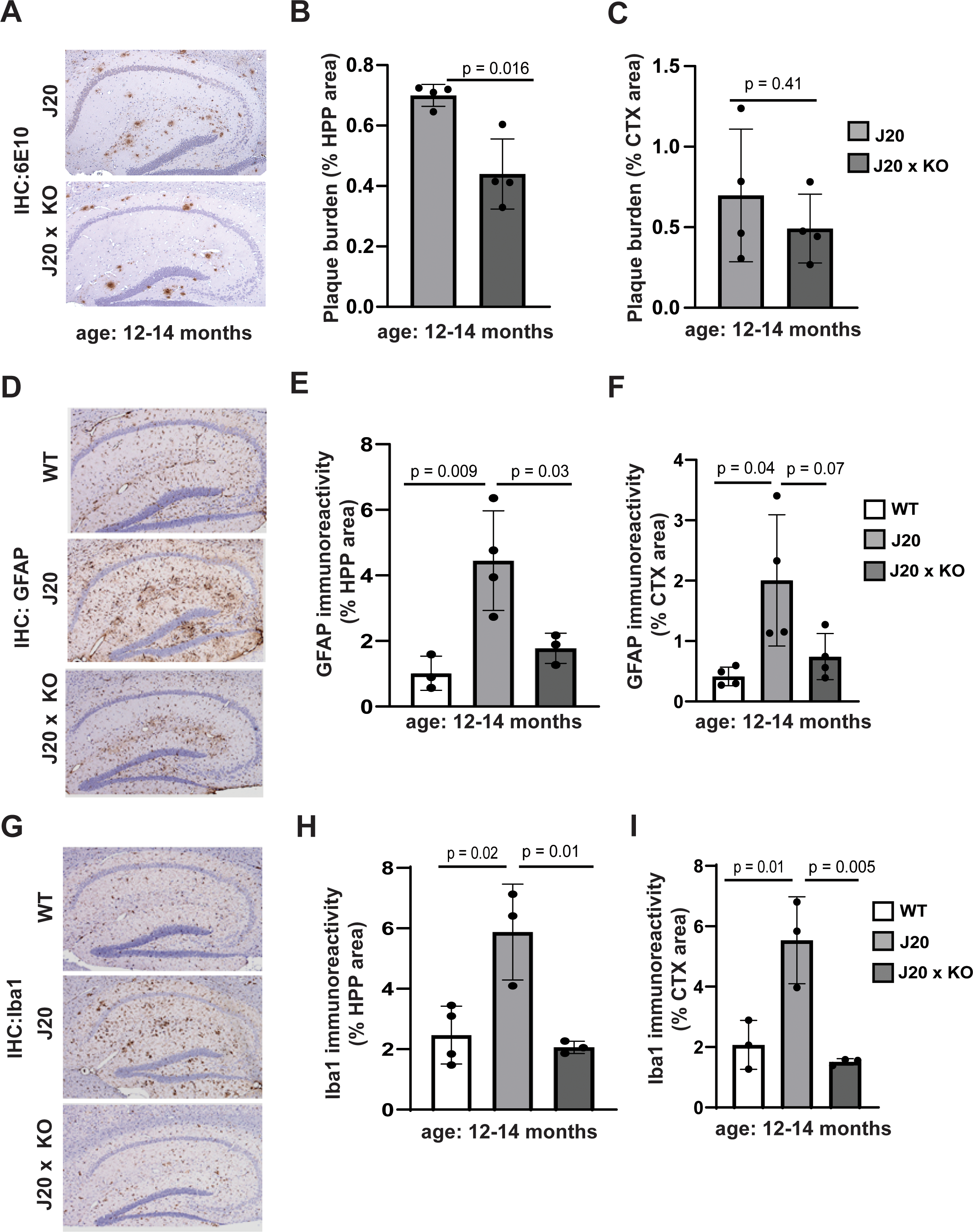
Effect of ADAP1/CentA1 KO on the histopathological hallmarks of AD in the J20 mice. ***A***, Representative images of amyloid plaque burden in the hippocampus in paraffin-embedded whole brain sections from hAPP-J20 and hAPP-J20 x ADAP1/ CentA1 KO mice. ***B***, Graph shows significantly reduced plaque burden in the hippocampus of J20 x KO mice compared to J20 mice (p=0.016). ***C***, Graph shows reduction in plaque burden in the cortex of J20 mice on ADAP1/ CentA1 KO background (p=0.41); however the difference did not reach statistical significance. ***D***, Representative images of GFAP immunoreactivity in the hippocampus in paraffin-embedded whole brain sections from WT, hAPP-J20 and hAPP-J20 x ADAP1/ CentA1 KO mice. ***E***, Graph shows significantly increased astrogliosis in the hippocampus of J20 mice compared to control mice and J20 mice on ADAP1/ CentA1 KO background. ***F***, Graph shows significantly increased astrogliosis in the cortex of J20 mice compared to control mice. ADAP1/ CentA1 KO reduced the astrogliosis, however the effect did not reach significance. ***G***, Representative images of Iba-1 immunoreactivity in the hippocampus in paraffin-embedded whole brain sections from WT, hAPP-J20 and hAPP-J20 x ADAP1/ CentA1 KO mice. ***H-I***, Graphs show extensive microgliosis in the hippocampus and cortex of J20 mice compared to WT controls; this effect is rescued in the J20 x ADAP1/CentA1 KO mice.

### Lack of ADAP1/CentA1 ameliorates neuroinflammation in the J20 mice

Detailed immunohistochemical analysis of 12-14 months hAPP-J20 mice showed higher astrogliosis (**FIG4. D-F)** and microgliosis (**FIG4. G-I)** compared with hAPP-J20 x CentA1 KO mice. Robust reactive gliosis was seen in the hippocampal CA1-CA3 region, periventricular areas, frontal cortex, and cerebellum of hAPP-J20 mice compared to hAPP-J20 x CentA1 KO mice. Quantitation of GFAP immunoreactivity indicated a substantial increase in astrocytic activation in the hippocampus and cortex of AD model mice compared to wild type littermates, that was significantly reduced in hAPP-J20 x CentA1 KO hippocampus (**FIG4. E;** WT: 1.018±0.26%; n=4; J20: 5.02±0.84%; n=4; J20 x KO: 2.5±0.8%: n=4; WT vs. J20: p=0.009; WT vs. J20 x KO: p=0.67; J20 vs. J20 x KO: p=0.03; S.E.M; one-way ANOVA followed by Tukey’s multiple comparisons test). Also, in the cortex, the hAPP-J20 x CentA1 KO mice had a trend toward lower astrogliosis than hAPP-J20 (**FIG4. F;** WT: 0.41±0.07%; n=4; J20: 2.00±0.54%; n=4; J20 x KO: 0.7±0.19%: n=4; WT vs. J20: p=0.04; WT vs. J20 x KO: p=0.9; J20 vs. J20 x KO: p=0.07; S.E.M; one-way ANOVA followed by Tukey’s multiple comparisons test). Widespread reactive microgliosis was also observed in the brain of the AD model mice, especially in and around the hippocampus (**FIG4. G-I)**. Quantification of Iba-1 immunoreactivity showed a robust increase in the expression of this microglia marker in the hippocampus of J20 mice (**FIG4. G and H)** compared with control mice and J20 mice on CenA1 KO background (**FIG4. H;** WT: 2.80±0.7%; n=4; J20: 5.87±1.25%; n=3; J20 x KO: 2.0±0.19%: n=3; WT vs. J20: p=0.02; WT vs. J20 x KO: p=0.68; J20 vs. J20 x KO: p=0.01; S.E.M; one-way ANOVA followed by Tukey’s multiple comparisons test). Similarly, extensive Iba-1 immunoreactivity was observed in the cortex of J20 mice (**FIG4. G and H)** that was significantly reduced by lack of ADAP-1/CentA1 (**FIG4. I;** WT: 2.0±0.7%; n=3; J20: 5.53±1.21%; n=3; J20 x KO: 1.5±0.09%: n=3; WT vs. J20: p=0.01; WT vs. J20 x KO: p=0.76; J20 vs. J20 x KO: p=0.005; S.E.M; one-way ANOVA followed by Tukey’s multiple comparisons test).

### Transcriptome profiling of J20 mice on ADAP1/CentA1 KO background

To gain insights into the molecular mechanisms that mediate by which ADAP1/CentA1 influence Alzheimer’s disease development and progression, we employed the NanoString nCounter platform and evaluated the expression profile of genes associated with neurodegeneration, neuroinflammation, and aging in our cohorts. RNA was isolated from the brains of WT, J20 and J20 x KO mice. We analyzed differentially expressed genes (DEGs; **Tables 1-3**) and fundamental themes of neurodegeneration (**Tables 4-6**) between genotypes (J20 vs. WT; J20xKO vs. WT; J20xKO vs. J20). Heatmap analysis was employed to depict the hierarchical clustering of DEGs, with downregulated genes being depicted in yellow while upregulated genes are depicted in blue **(FIG. 5-7).**

**FIG. 5:**
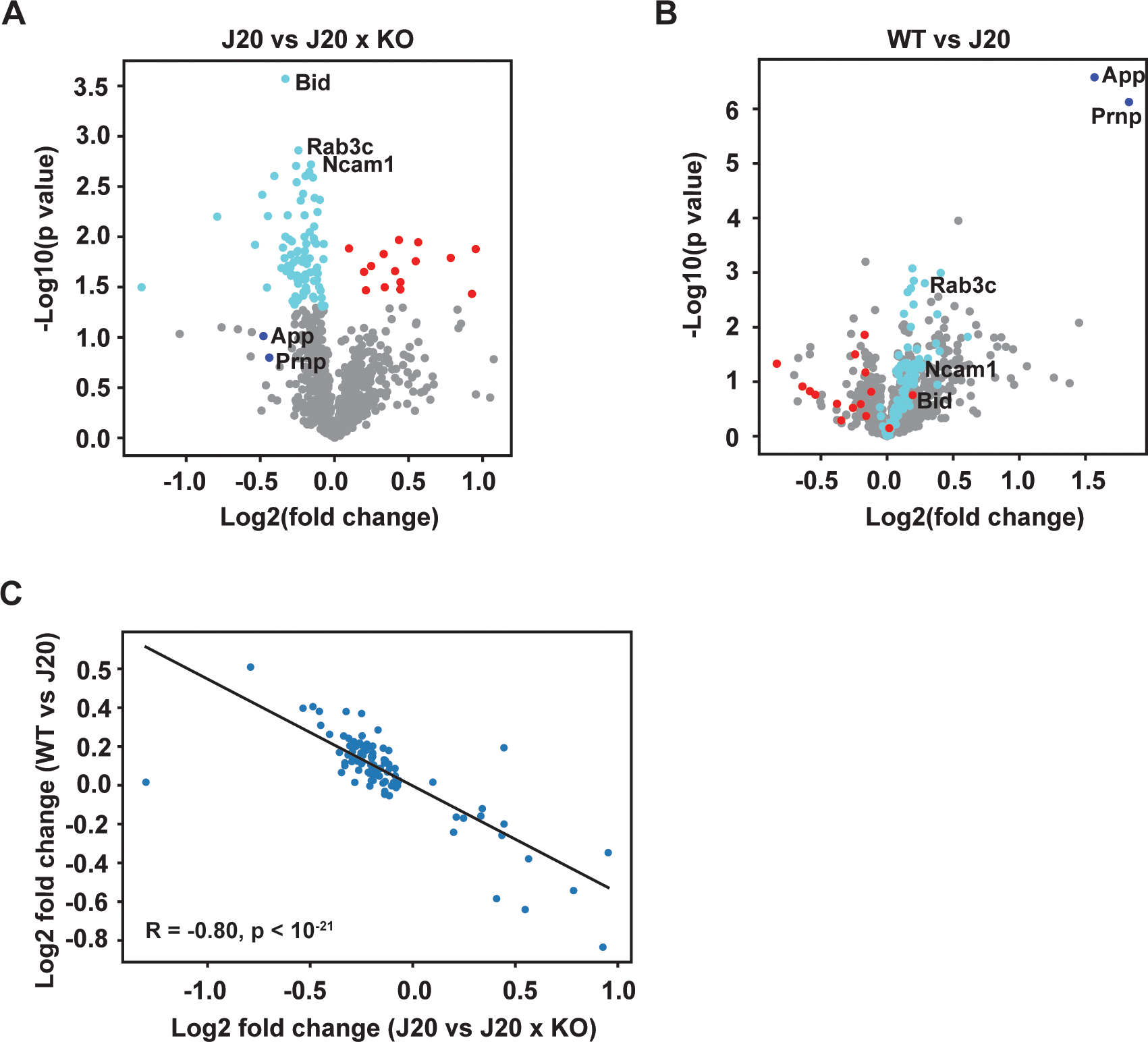
Differentially expressed genes between hAPP-J20 and WT mice identified with NanoString nCounter profiling. Heatmaps depict the hierarchical clustering of differentially expressed genes (DEGs) between genotypes, of which 17 (**A**) were significantly downregulated and 74 (**B**) were significantly upregulated in the forebrain of hAPP-J20 mice compared to their wild type (WT) littermates. The list of DEGs and statistical analysis are presented in **Table 1**. For the association of the DEGs with fundamental themes of neurodegeneration, see **Table 4**.

**FIG. 6:** Differentially expressed genes between hAPP-J20 x CentA1 KO and WT mice identified with NanoString nCounter profiling. Heatmaps depict the hierarchical clustering of differentially expressed genes (DEGs) between genotypes, of which 19 (**A**) were significantly downregulated and 29 (**B**) were significantly upregulated in the forebrain of hAPP-J20 x ADAP1/ CentA1 KO mice compared to their wild type (WT) littermates. The list of the genes and statistical analysis are presented in **Table 2**. For the association of the identified DEGs with fundamental themes of neurodegeneration, see **Table 5**.

**FIG. 7:** Differentially expressed genes between hAPP-J20 and hAPP-J20 x CentA1 KO mice identified with NanoString nCounter profiling. Heatmaps depict the hierarchical clustering of differentially expressed genes (DEGs) between genotypes, of which 73 (**A**) were significantly downregulated and 14 (**B**) were significantly upregulated in the forebrain of hAPP-J20 x ADAP1/ CentA1 KO mice compared to their wild type (WT) littermates. The list of the genes and statistical analysis are presented in **Table 3**. For the association of the identified DEGs with fundamental themes of neurodegeneration, see **Table 6**.

Our data shows that the fundamental themes of neurodegeneration most affected by the J20 genotype are metabolism, compartmentalization, structural integrity, neurotransmission, neuroplasticity, development, and aging (**FIG. 8; Table 7**). Among these genes, we further analyzed Bid, because Bid is known for XXX. We validated the changes in Bid expression by quantitative RT-PCR (qRT-PCR) analysis **(FIG. 9A)**, which was performed on RNA isolated from the corresponding frozen tissue specimens used for NanoString nCounter analysis ^38,39^. Similar changes were observed at Bid individual gene level, showing statistical significance in the qRT-PCR analysis (WT: 0.80±0.12; n=4; J20: 3.45±0.86; n=6; J20 x KO: 0.59±0.15; n=5; WT vs. J20: p=0.03; J20 vs. J20 x KO: p=0.01; S.E.M; one-way ANOVA followed by Tukey’s multiple comparisons test). Next, we validated the NanoString and qRT-PCR results using biochemical analysis of the corresponding frozen tissue specimens. Western blotting analysis of Bid protein level in the hippocampus **(FIG. 9B)** showed reduced expression of Bid in J20 mice on KO background compared to both WT and J20 genotypes.

**FIG. 8:** Fundamental themes of neurodegeneration most affected by genotype. ***A***, Graph shows association of DEGs with themes of neurodegeneration in the forebrain of hAPP-J20 vs. WT mice. The most affected pathways are neuroplasticity, development, and aging; neuroinflammation; neurotransmission; metabolism; compartmentalization, and structural integrity. ***B***, Graph shows association of DEGs with themes of neurodegeneration in hAPP-J20x ADAP1 KO vs. WT mice. The most affected pathways are neuroplasticity, development, and aging; neuroinflammation; neurotransmission; metabolism; compartmentalization, and structural integrity. ***C***, Graph shows association of DEGs with themes of neurodegeneration in hAPP-J20x ADAP1 KO vs. hAPP-J20 mice. The most affected pathways are neuroplasticity, development, and aging; neurotransmission; metabolism; compartmentalization, and structural integrity.

**FIG. 9.** Quantitative RT-PCR and western blotting to validate a subset of DEGs. ***A***, Validation of the changes in *Bid* at individual gene level expression using quantitative RT-PCR (qRT-PCR). Similar to bulk gene expression analysis, *Bid* was significantly increased in the forebrain of AD mice compared to WT and hAPP-J20 mice on CentA1 null background. ***B***, Biochemical analysis of the level of Bid protein in the forebrain indicates reduced expression of Bid in hAPP-J20 x CentA1 KO mice compared to both WT and hAPP-J20 genotypes.

## DISCUSSION

In the present study, we performed a detailed and multimodal analysis of the involvement of ADAP1/CentA1 in the pathophysiology of Alzheimer’s disease (AD). Previous research on the roles of ADAP1 in brain health and disease implicated this protein in AD pathology^11,23^ and identified ADAP1 as a negative regulator of dendritic spine density, sLTP and learning and memory^29^. We therefore hypothesized that in mice expressing hAPP on *Centa1* null background, key aspects of AD (learning and memory deficit, synapse loss, amyloid plaque burden and neuroinflammation/gliosis) will be reduced or rescued. For our experiments, transgenic mice expressing hAPP with the Swedish and Indiana FAD mutations (line J20; hAPPJ20 mice)^30^ were crossed with *Centa1* null (ADAP1/CentA1 KO) mice^29^. In an initial morphological analysis, we did not observe substantial differences in gross brain morphology of ADAP1-deficient J20 mice compared to WT/NTG and hAPP J20 littermates (**FIG.1A**). Using immunoblotting we also validated the lack of ADAP1 expression and the presence of transgene (hAPP) in the hippocampus of 6 months old (**FIG.1B**) and 12–14-month-old J20xKO mice (**FIG.1C**). The brain regions affected the earliest in AD pathogenesis in both humans and animal models are the areas associated with spatial navigation and orientation^3,40,41^. In the J20 mice, deficits in spatial memory and learning are evident at 5-7 months of age^33,42^. Consistent with previous studies^42,43^, the performance of the hAPP-J20 mice was significantly impaired in the acquisition phase of the Morris Water Maze tests (MWM), with the mice needing nearly double the number of trials to reach criterion compared to their WT and KO littermates (**FIG.2**). Interestingly, this effect was rescued by the deletion of ADAP1/CentA1, suggesting that ADPA1/CentA1 plays a critical role in APP-induced memory loss. Given that spatial learning and memory performance are thought to correlate with changes in dendritic spine number and morphology^44^ ^45–47^, we also tested the effects of *Centa1* deletion on dendritic spine loss caused by cerebral Aβ amyloidosis in hAPP-J20 mice. Previous reports indicated a spatially restricted loss of synapses and dendritic spines in the hippocampal CA1 neurons of J20 mice^48–50^. Therefore, we analyzed spine density in the hippocampal stratum lacunosum moleculare (SLM; receives direct presynaptic inputs from the entorhinal cortex) and in the stratum radiatum layer (SR; receives inputs from CA3). We found that in SLM, spine density of CA1 neurons of J20 mice was significantly reduced compared to NTG littermates and this loss of spines was rescued by lack of ADAP1 (**FIG.3A**). However, spine density was significantly higher in the SR layer of J20 mice, with no effect of CentA1-defficit (**FIG.3B**). This may be due to subcellular domain-specific CentA1 expression. Spatially restricted cell morphology changes in neurons of J20 mice have been reported recently; these changes included mitochondrial fragmentation and loss in the apical tufts of CA1 pyramidal neurons, coinciding with the earliest spine elimination *in vivo*^50^. This mechanism is supported by previous studies showing that ADAP1/CentA1 accumulation in mitochondria destabilizes mitochondria via Ca^2+^-induced opening of the permeability transition pore (PTP)^11,51^. These results suggest that Centaurin-α1 is a key factor in a novel mechanism regulating dendritic spine loss and neural dysfunction that ultimately leads to the behavioral abnormalities associated with the AD-phenotype.

One important hallmark of AD is the formation of amyloid plaques throughout the brain. ADAP1/CentA1 is highly expressed in the human AD brain, particularly around senile plaques^27^. Age-dependent expression of Aβ, followed by senile plaque deposition at later stages in the hippocampus, has been extensively described in the hAPP-J20 model^33^. Therefore, we evaluated whether the lack of ADAP1/CentA1 might impact plaque formation in the J20 mice. We found significantly reduced plaque burden in the hippocampus of J20 x KO mice, while cortical plaque formation trended toward a reduced level (**FIG.4A-C**). Potential signaling pathways mediating the reduction of plaque burden in J20 mice lacking CentA1 include the small GTPases Ras and Arf6, as AD-associated downstream effectors of CentA1 signaling^25,26,52^. Ras signaling has been shown to trigger APP expression, and in turn, APP and Aβ activate the Ras-ERK signaling to promote dendritic spine loss and neurodegeneration^14,53–55^. Increased CentA1 expression as AD progresses might exacerbate this feedback loop. Indeed our previous studies show an Aβ-dependent activation of the CentA1-Ras-ERK-Elk1 pathway, while CentA1-down regulation in the hippocampus reduces Ras activity^11,29^. On the other hand, Arf6 activity increases in the brain lacking CentA1^29^ and Arf6 has been shown to regulate APP processing via sorting BACE1 to early endosomes within the somatodendritic compartment of polarized hippocampal neurons^56^. In this model, Arf6 controls the access of BACE1 to its substrate APP; consequently, increasing Arf6 activity results in increased amyloidogenic processing of APP and enhanced Aβ generation. The mechanism by which these two pathways crosstalk to control Aβ production and secretion in the absence of CentA1 are unknown.

Neuroinflammation is another neuropathological hallmark of AD. Reactive microgliosis and astrocytosis produce inflammatory cytokines and chemokines, exacerbating neuronal dysfunctions and promoting neurodegeneration^57,58^. Recent studies have shown the involvement of pro-inflammatory, neurotoxic A1 reactive astrocytes in the initiation and progression of AD^57,59^.

Comprehensive analysis of microglia states in different stages of AD, revealed that reactive microgliosis strikingly follows the spatio-temporal progression of the disease and requires the presence of both amyloid and tau pathology^39^. Although ADAP1/CentA1 was previously thought to be a neuronal-restricted factor, recent studies have indicated that astrocytes^60^ and immune cells^61^ also express this protein. Therefore, we evaluated whether lack of ADAP1/CentA1 might impact neuroinflammation in the J20 mice. We found significantly reduced reactive astrogliosis (as indicated by GFAP-immunoreactivity) in the hippocampus of J20 x KO mice, while cortical astrogliosis was slightly reduced without reaching statistical significance (**FIG.4D-F**). Iba-1 immunoreactivity was significantly reduced in J20 mice on KO background in both the hippocampus and the cortex, suggesting that ADAP1/CentA1 signals to microgliosis (**FIG.4G-I**).

Recent transcriptome and epigenetic studies revealed the downregulation of neuronal functions and upregulation of innate immune responses with disease progression, indicating that altered neuron-glia interactions are key players in the pathophysiology of AD^5,62,63^. To gain further insights into the pathways driving AD-like neuropathology in the hAPP-J20 mice and to evaluate the role of ADAP1/CentA1 in this process, we performed whole tissue transcriptomics on the NanoString nCounter^®^ platform. We aimed to evaluate gene expression profiles and identify novel changes in the fundamental themes of neurodegeneration potentially associated with lack of ADAP1/CentA1. Hence, we compared neurodegeneration-associated gene expression and biological processes in the hippocampal tissue of hAPP-J20 mice, hAPP-J20 mice on CentA1 KO background and their NTG/WT litter mates (**FIG.5-8**). As expected in bulk brain tissue RNA transcriptomics, the expression signals of neurons and oligodendrocytes were the most abundant (**FIG 8**). The largest difference in pathology-responsive transcriptional signatures was observed between the hAPP-J20 and NTG/WT hippocampi (**FIG.5 and 8A**). We found that hAPP-J20 mice show upregulation of genes associated with oxidative stress, unfolded protein response, reactive microgliosis, and dysfunctional neuronal compartmentalization and structural integrity, particularly myelination **(Tables 1, 4 and 7)**. Notably, the number of DEGs between hAPP-J20 mice on CentA1 KO background and the NTG/WT was lower (**FIG.6 and 8B**) and the most affected biological processes included the pathways of compartmentalization, structural integrity, neuroinflammation, neuroplasticity, development, and aging **(Tables 2, 5 and 7)**. Importantly, the perturbations in gene expression by the hAPP-J20 genotype were largely rescued by the deletion of ADAP1/CentA1 (**FIG.7 and 8C**). The altered biological processes include metabolism; neurotransmission; compartmentalization and structural integrity; neuroplasticity, development, and aging. Transcriptional signatures of neuronal death and survival, inflammation, and myelination were among the most altered pathways **(Tables 3, 6 and 7)**. We found that one of the top DEGs was Bid (**FIG.7B**), upregulated in the hAPP-J20 mice, in parallel with downregulation of the anti-apoptotic factor, BCL2L1 (**FIG.7A**). As a pro-apoptotic factor in the mitochondrial pathway of apoptosis^64^, Bid has been shown to be involved in caspase activation associated with AD^65^. Given the mitochondrial localization and role of ADAP1/CentA1^23,51^, we further validated the upregulation of Bid (**FIG. 9**) in the hAPP-J20 hippocampi and rescue by lack of ADAP1 using qPCR and also western blot analysis. Given that many transcriptional alterations are cell-type and brain region-specific^66^ and expression of ADAP1/CentA1 varies between brain areas^19,28^, an important future direction is the use of single-cell spatial transcriptomics to evaluate the effect of CentA1 KO on the progression of neuropathological hallmarks in AD-vulnerable areas. Combining anatomical information with cell-specific transcriptional alterations, could lead to identification of shared signaling pathways across major cell types and identify novel therapeutic targets.

Overall, we identified ADAP1/CentA1 as a potential mediator of Alzheimer’s disease (AD) pathogenesis. Our results highlight the involvement of myelination-, neuroinflammation-, cell death- and survival-related processes in pathophysiology of AD and provide evidence for a neuroprotective outcome of lowering the level of ADAP1/CentA1 in the brain.

## ACKNOWLEAGEMENTS

Authors thank Minida Dowdy and the ARC at MPFI staff for outstanding and compassionate animal care; Boram Yu (MPFI), Erin Suellentrop and Elizabeth Brennan (ECU) for technical support; to Paramita Chakrabarty (University of Florida) for expert advice on brain pathology analysis; to Yasuda, Stackman and Szatmari lab members for constructive discussions and review of the manuscript. This work was supported by generous funds from the BrightFocus Foundation, Community Foundation of the Palm Beaches, NINDS, Max Planck Foundation, ECU Startup funds and ECU URCA awards.

